# Shape that matters: Yolk geometry spatially modulates developing vascular networks within chick chorioallantoic membrane

**DOI:** 10.1101/2024.07.18.604146

**Authors:** Prasanna Padmanaban, Dirk Wanders, Oryan Karni Katovich, Nasim Salehi-Nik, Shulamit Levenberg, Jeroen Rouwkema

## Abstract

Controlling the multiscale organization of vasculature within diverse geometries is essential for shaping tissue-specific and organ-specific architectures. Nevertheless, how geometrical characteristics of surrounding tissues influence vessel morphology and blood flow remains unclear. Where the regulation of vascular organization by mechanical signals associated with fluid flow is well known, this study postulates that the organization of developing vasculature can also be regulated by mechanical signals connected to the confinement and thus the deformation of surrounding tissues. To test the Shape-Induced Vascular Adaptation (SIVA) concept, fertilized chicken egg contents containing developing vasculature were cultured within engineered eggshell platforms of different shapes. Our findings demonstrate that the vascularized chick chorioallantoic membrane (CAM) adapts to the shape of engineered eggshell, long before reaching its boundaries. This adaptation affects the organization of the vascular network within the CAM, affecting parameters such as vessel area, branching, orientation, length, diameter and endpoints. Specifically, we observed that sharp corners in the engineered eggshell led to more elongated vascular structures. To further explore the dynamic nature of this phenomenon, a proof-of-concept experiment was performed using a shape-shifting engineered eggshell that deforms the egg content from circle to square shape. Using this shape-shifting prototype, we observed a direct effect of eggshell deformation on the vessel morphology and flow dynamics in a time-dependent manner. Overall, our ex-ovo experimental platform provides a unique opportunity to study how mechanical stimuli such as shape influence the spatial and temporal organization of developing vascularized tissues. By subjecting these tissues to various static and dynamic conditions, we induced both local and global changes in their organization. This class of perturbation provides us with an additional tool which can be used for shaping vascular organization within developing tissues and to engineer tissues with geometrically tunable vessel structures.

## 1. Introduction

Mechanical forces such as shear stress^[1,2]^, circumferential strain^[3]^ due to pulsatile blood flow, compression forces^[4]^, and tissue rigidity^[5–7]^ are crucial for vascular tissue development and vascular network architecture. These signals that are connected to the flow of blood through the vasculature regulates sprouting and remodeling processes, resulting in blood vessels with varying diameters, lengths, and branching patterns. The effect of these natural signals are well studied. However, artificial in vitro environments that are used for instance in tissue engineering, offer the opportunity to also control and perturb other factors, such as tissue geometry. Previous research has shown that in an engineering setting changing the tissue geometry has an effect on vascular formation and organization^[8,9]^. However, these processes and the mechanisms behind it are less well studied.

Morphological transitions such as the formation of tubes, lumens, complex curved loops, and branching structures are observed during embryonic development to shape vasculogenesis, angiogenic sprouting and vascular network organization^[10–20]^. Recent studies show that mechanical forces like pressure^[21,22]^, compression^[4,23,24]^, and stretch are crucial in molding the vascularized tissue shapes. Recently, we have demonstrated that the development and organization of vascular networks during embryogenesis are adaptable to flow-induced shear stresses and can be altered by externally induced flows^[25,26]^. In engineered microtissues, local mechanical strain emerging from compaction and cellular traction forces on the extracellular environment induce tissue deformation, leading to changes in topology and organization of vasculature^[8]^. Furthermore, observations from plant biology indicate that branched tree structures and the overall shapes of fruits can be altered using external molds ^[34,35]^. Inspired by these findings, we aim to explore methods to actively guide the development and organization of vascular networks using external shapes to exert localized control over strain and stresses.

In this manuscript, we present an innovative technique called shape-induced vascular adaptation (SIVA). This technique involves culturing of developing vascularized chick chorioallantoic membrane (CAM) tissues within engineered eggshell platforms of different shapes **(Figure 1)**. Engineered eggshell platforms are simple, cost-effective and powerful tools to systematically investigate and manipulate the vasculature of chick embryo over the course of development. We fabricate engineered eggshell platforms using a polydimethylsiloxane (PDMS) based molding technique to achieve differently shaped eggshell including circle, square, triangle and star designs. These eggshells can accommodate entire chick CAM specimens, which include diverse multiscale vasculature networks **(Figure 2)**. Using this methodology, we observed significant differences in the vasculature organization of CAM tissue when comparing the different shapes, including changes in vessel area percentage, lacunarity, average vessel length, diameter, branching, orientation and end points **(Figure 3-5)**. To further explore the dynamic nature of this phenomenon, we engineered a shape-shifting prototype that could transform eggshell from circle to square using pressure-driven pneumatic actuators **(Supplementary Video 3)**. By applying compression actuation, we introduced changes in the vascular organization within CAM tissue and closely examined how this impacted the vascular organization in multiple regions of the CAM tissue **(Figure 6)**. Overall, this study shows that mechanical cues connected to the shape of a tissue can influence vascular organization within developing embryo tissues. Our findings serve as a basis for further research into the signaling mechanisms of shape-induced engineering of vascularized tissues.

**Figure 1.**
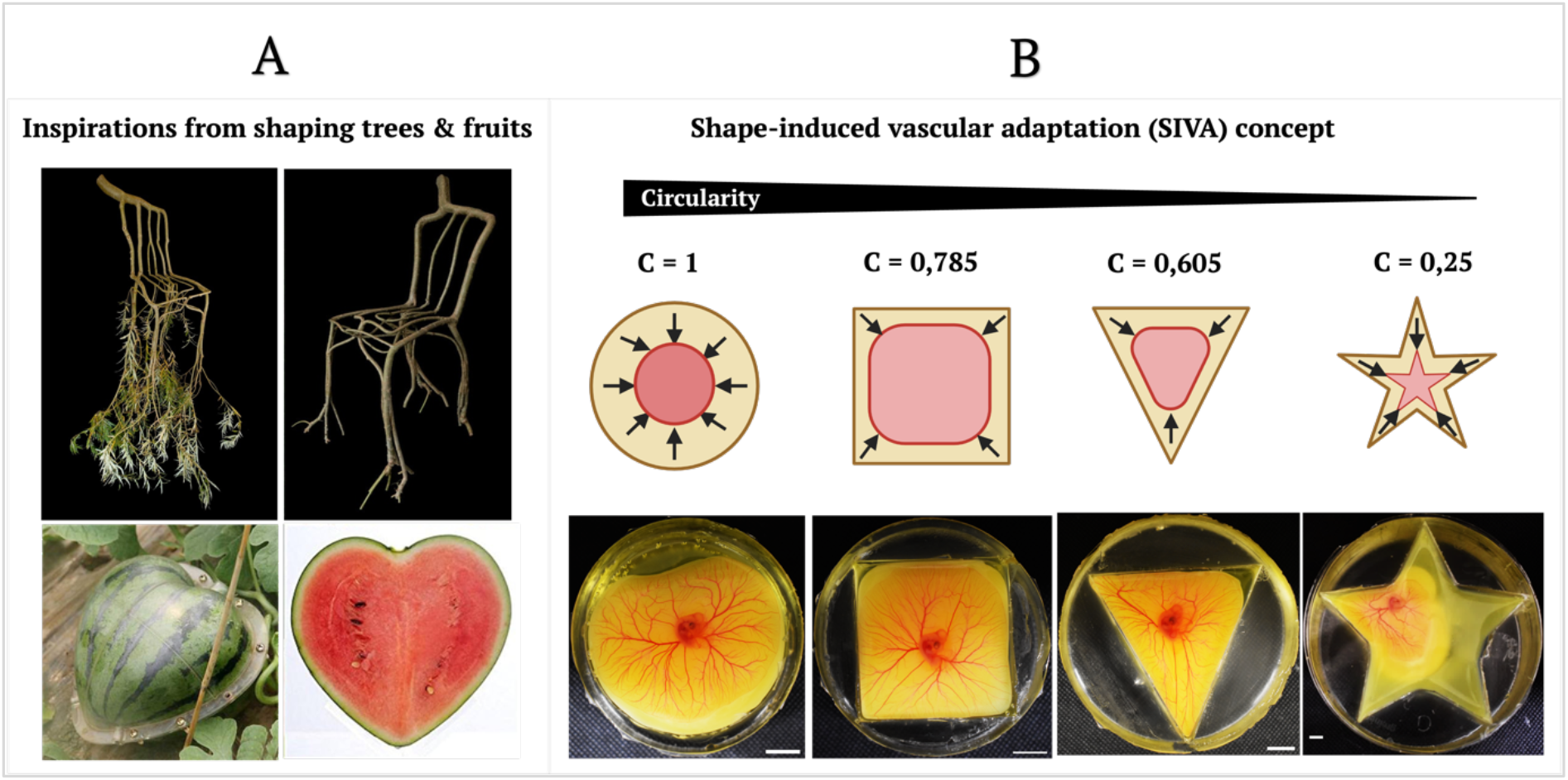
Shape-induced vascular adaptation concept. (A) The top panel illustrates the “shaping trees” concept developed by the Full-Grown company, where growing trees are directed around molds to create a chair shape. The bottom panel shows plastic molds wrapped around growing watermelons, causing them to take on a heart shape. (B) This panel demonstrates how chick yolk adapts to the shape of different engineered eggshell platforms, leading to changes in the organization of vascular networks. Scale bar represents 10mm.

**Figure 2.**
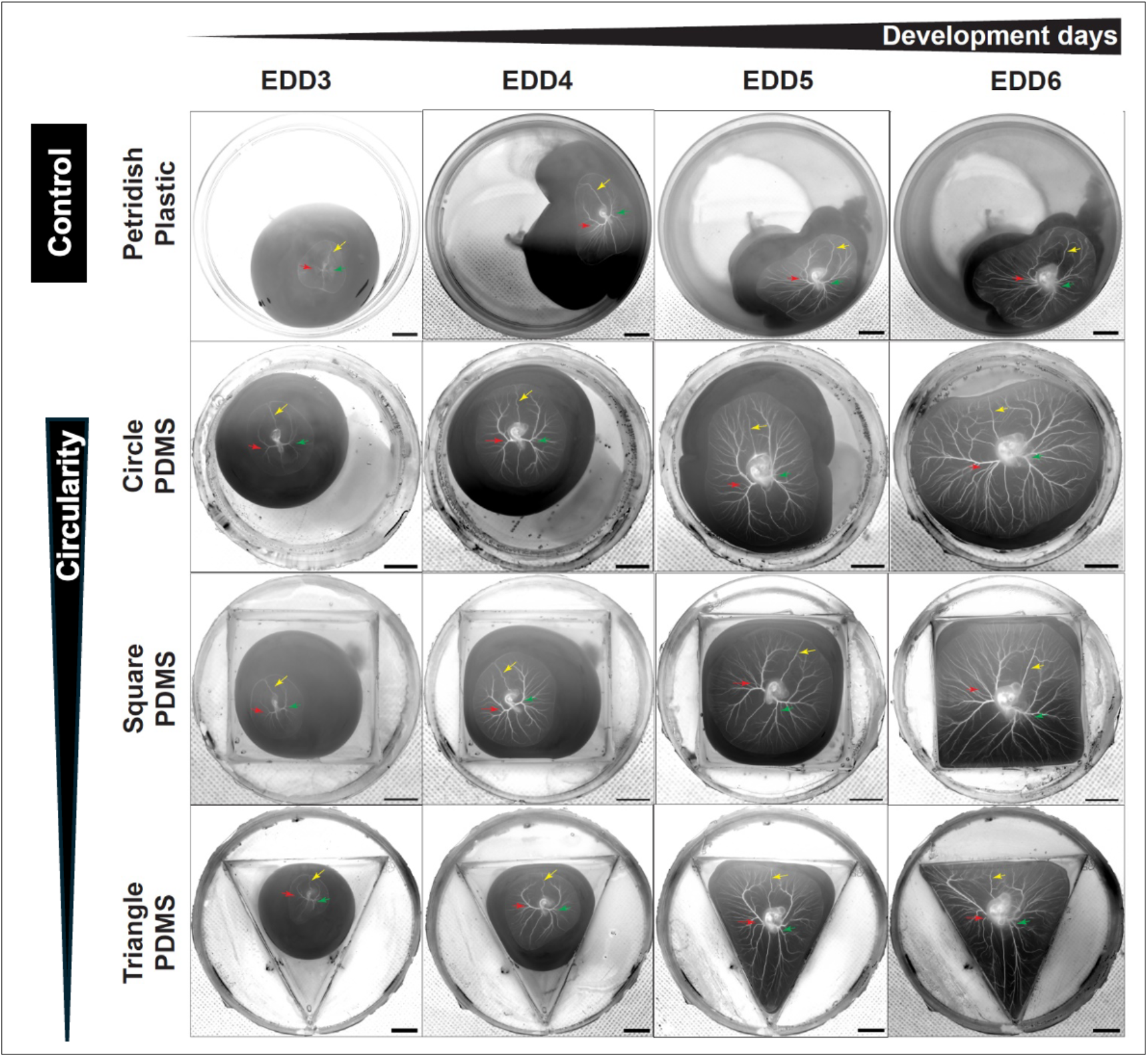
Spatiotemporal organization of chick vasculature grown in shaped culture platform. Snapshots of the entire vasculature of developing chick embryos cultured within differently shaped, engineered egg containers from embryonic development days (EDD) 3 to 6. The figure showcases the evolution of yolk shape and their respective CAM vasculature, demonstrating that organization is influenced by the shape of culture geometry. Red arrows indicate the left side symmetry of first main hierarchical branching vessels arising from the embryo, green arrows indicate right side symmetry of first main hierarchical branching vessels arising from the embryo expanding towards the surrounding yolk sac membrane of CAM tissue and yellow arrows indicate the straight vessel connecting the marginal vessel to the embryo respectively. **First row** (plastic Petri dish) shows an increasing number of branching vessels, along with an increase in vessel length, and tortuosity of vessels present of the left side. **Second row** (circle geometry) exhibits well maintained circularity of yolk and CAM with arc shaped branching vessels with excessive sprouting over the development. **Third row** (square geometry) portrays elongated branching vessels over development and **fourth row** (triangle geometry) represents elongated vessels at the bottom side and increased sprouting at the top side. Orientation and directionality of vascular structures are strongly influenced by sharp corners. Scale bar represents 10mm.

**Figure 3.**
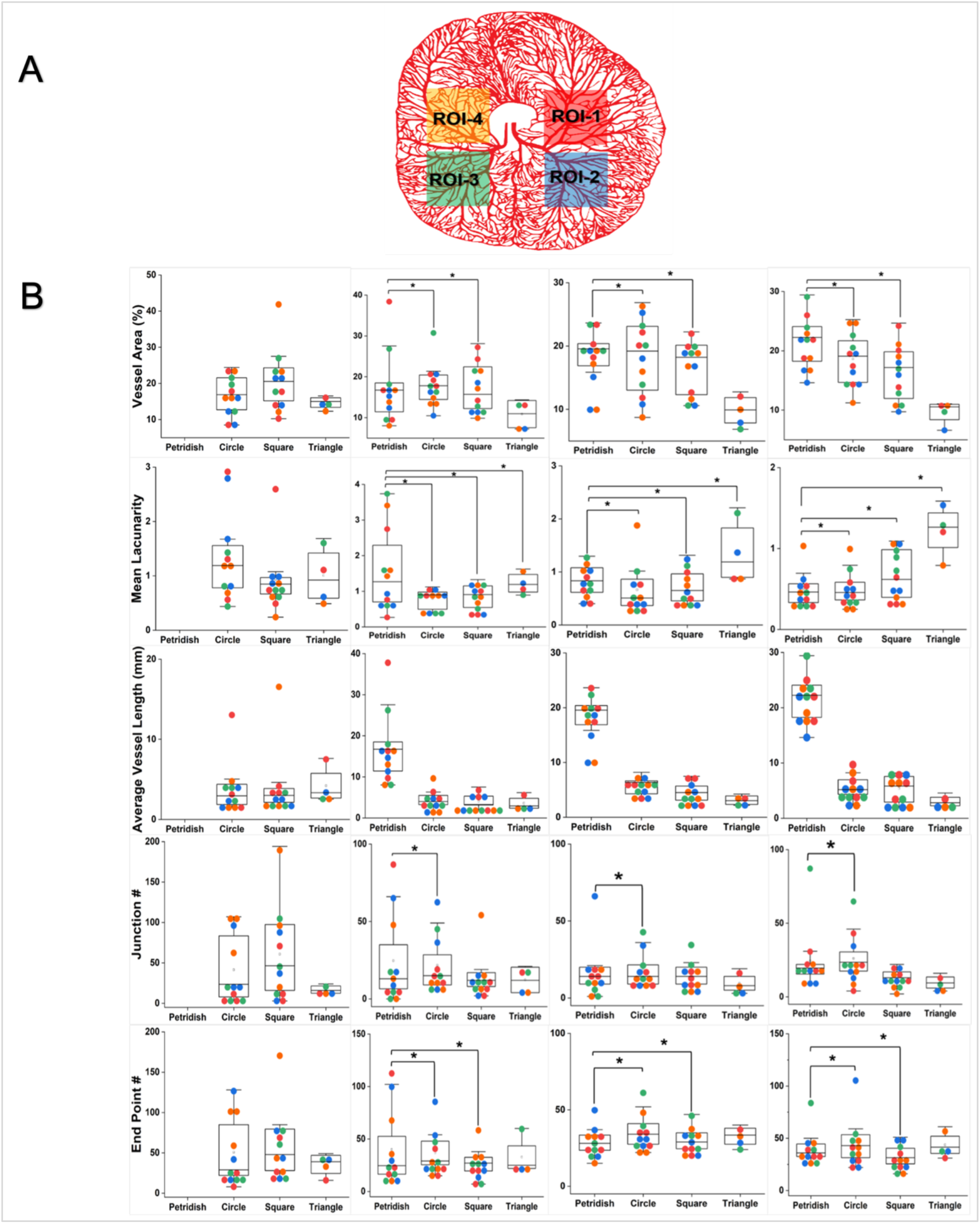
Morphometrics analysis and influence of culture platform shapes on vascular structures. **(A)** represents the schematic mask of chick vascular network created using Adobe illustrator with highlighted regions of interest (ROI) represented for the quantitative analysis. A region of 500 × 500 pixels corresponding to each ROIs is used for morphometric evaluation. **(B)** shows the graphical representation of quantitative data analysis results. Measured values are represented as mean ± SD along with individual data points compared within 4 different ROIs from different culture platform and development days as indicated. Color coded labels were used for ROI1(red), ROI2(blue), ROI3(green) and ROI4(orange) respectively. Measured values are calculated from 3 individual samples (n=3) for all shapes, except triangle shape where only one individual sample is considered. Note that data points for Petri dish for EDD3 were excluded because insufficient vascular structures were observed. Statistical significance test was performed using OriginPro V.9.85 performing one way ANOVA and differences were considered statistically significant with P<0.05

## 2. Experimental section

### 2.1. Ethics statement

In accordance with the Dutch animal care guidelines, obtaining Institutional Animal Care and Use Committee (IACUC) approval for experiments involving chicken embryos is not mandated unless there is an anticipation of hatching. Additionally, IACUC approval is only requisite for experiments involving chick embryos at or beyond EDD14 of development. The embryos utilized in this research were all in the early stages of development, falling between EDD3 and EDD6. The fertilized chicken eggs employed in this study were procured from authorized poultry egg farms in the Netherlands.

### 2.2. Chick embryo culture and transfer to engineered egg container

Chicken embryos were grown as reported earlier^[25]^. Dekalb white fertilized chicken eggs were obtained from Het Anker BV in Ochten, The Netherlands, and stored at 12 °C. A Day before egg incubation, a modified incubator was prewarmed to 38 °C with 65% humidity, which was maintained throughout incubation. In the initial 3-day incubation period, the eggs were rotated every two hours for 15 seconds to prevent embryo adhesion to the eggshell. 3 days post the start of incubation, a small ∼2 mm diameter hole was created in the eggshell with fine tweezers, and an 18G syringe microneedle with a plastic syringe was used to withdraw 3 mL of albumen to protect the yolk and embryo vasculature during cracking as shown in **Supplementary Figure S1**. Carefully, the withdrawn albumen was introduced into the engineered eggshell container to minimize potential disturbance of the yolk during transfer. After this, the whole remaining egg content, including the chicken embryo and the CAM containing multi-sized vascular networks, was transferred to the engineered eggshell containers in a sterile environment.

### 2.3. Shaped egg containers and shape-shifting prototype fabrication

As reported earlier, the egg containers were made of PDMS (Sylgard 184 silicone elastomer, Dow corning)^[25]^. The PDMS base was mixed at 10:1 (w/w) with the curing agent and poured into a laser cut polymethyl methacrylate (PMMA) block of different shapes as shown in **Supplementary Figure S2**. Shapes were standardized to have a cross-sectional surface area of 5 cm2 and a height of 3 cm. The mold was then placed in a desiccator to remove potential air bubbles, and the PDMS was cured overnight in a 65 ºC dry oven. Later, the cured PDMS and PMMA mold were carefully detached after washing with soap solution for easy removal. A shape shifting prototype was fabricated by assembling PMMA blocks with pneumatic cylindrical actuators driven by directly acting 2-way solenoid valve operating pressure ranges between 0.15 and 0.8MPa, performing compression actuation.

### 2.4. Image acquisition

Whole vasculature snapshots of differently shaped egg containers and circulating red blood cells videos during shape-shifting actuation were captured using a combination of three cameras: Dino-Lite USB camera (AM4115 ZT 1.3MP), an HAYEAR digital microscope (16MP Industrial grade, 150× C-Mount Lens) and Nikon DSLR D3500 with 55mm Lens. The recorded image and video files were processed using the corresponding software: DinoXcope 2.0 for Mac and IC Measure software for Windows. Multiple frames extracted from the processed images were then imported and transformed into 8-bit grayscale images using Fiji software.

### 2.5. Image processing and data analysis

The detailed step-by-step analysis of vessel morphometrics, vessel diameter, and vessel orientation is explained in **Supplementary Figure S4**. Various vessel morphometric properties, such as vessel area percentage, lacunarity (void space), average vessel length, number of junctions/branches, and number of endpoints, were characterized using the Angiotool Version 0.6a Fiji plugin. First, raw images were converted into 8-bit grayscale images with selected regions of interest, each having a pixel value of 300 × 300. These images were then locally thresholded and imported into the Angiotool app for further analysis.

For the vessel diameter analysis, a few additional steps were incorporated. A Mexican hat filter with a radius value of 4.0 was applied to the 8-bit images, followed by an FFT bandpass filter with a large structures filter value of 100 pixels and a small structures filter value of 1 pixel, with a tolerance direction of 5%. The images were then smoothed, and local thickness was applied to plot the geometry onto the distance map. Finally, the diameter measurement plugin was used on the selected region of interest with a pixel value of 25 × 25.

As previously reported^[9]^, quantitative vessel orientation maps were generated using Matlab Version 2019b. Masks of the selected regions of interest were skeletonized and transformed into single vessel elements. These elements were classified based on the angle of their major length axis. The lengths of the elements were accumulated in 5º bins and plotted on a radial axis, where the distance from the origin represents the total length of all the binned elements, and the angle represents the orientation of the bin. The plots cover angles from -90º to 90º, reflecting the symmetry of the compartments along the y-axis.

## 3. Results and discussion

### 3.1. Shape-induced vascular remodeling and organization of chick CAM

Our objective was to evaluate the impact of shape on the vascular development and organization of chick chorioallantoic membrane cultured within differently shaped eggshell platforms. To accomplish this, we first engineered PDMS-based eggshell platforms of different shapes including circle, square, triangle and star shape as described in methods section 2.3 and **Supplementary Figure S2**. These shapes were inspired by previous in vitro designs that studied the effect of local geometry and topology on the development and orientation of sprouting vessels within fabricated microtissues^[8,9]^. Our transparent eggshell platforms can hold up to 60ml of fertilized chick egg contents including albumen and chick embryo containing multiscale vascular networks. This allowed us to image and compare the entire CAM vasculature across different shapes.

We observed significant differences in the vascular morphology and network topological organization when comparing the different shapes as shown in **Figure 2**. These differences were most prominent in the three main vessels, highlighted with colored arrows. These vessels are part of the embryo proper, as they extend from the embryo, either distributing oxygenated blood throughout the tissue or returning deoxygenated blood from the embryo to the CAM, where oxygenation happens. There is a clear asymmetry in vessel branching and tortuosity between the shapes, which appears to be influenced by how the yolk expands. This demonstrates that shape is an important regulator of vascular angiogenesis, remodeling and organization within the embryo over the course of development. Additionally, we observed that the diameter of the marginal vessels at the CAM boundary decreased as development progressed between shapes. However, the poor spatial resolution of these vessels made quantification very challenging. Notably, the results for the star shape morphometric analysis are excluded from **Figure 2** due to the yolk damage caused by asymmetric sharp corners leading to difficulties in long-term culturing. The **Supplementary Figure S3** and **Supplementary Video 1 and 2** demonstrate these effects and observed organizational changes within the star-shaped egg containers.

### 3.2. External shapes significantly influence vessel morphometrics, circularity and orientation

To compare the CAM vasculature development across different shapes, we captured time snapshots of entire vasculature from EDD3 to EDD6 as detailed in the methods section 2.4. Subsequently, we performed morphometric analysis using Angiotool. The results revealed an overall increasing trend in lacunarity and decreasing trends in vessel area, vessel length, junction number, and endpoint number in triangles and squares, compared to circles **(Figure 3B)**. Additionally, our findings, in conjunction with previously reported results confirm that shapes with more sharp corners, such as triangles and squares, lead to more vascular remodeling, when compared to circles.

Vessel diameter is an interesting morphological property for bioengineers to tune, particularly since fabrication techniques like 3D printing heavily rely on the diameter of the printing nozzle. We compared the left-side and right-side vascular symmetry of developing CAM and found significant differences in vessel diameters between the different shapes and control Petridish samples **(Figure 4C-F, Supplementary Figure S5)**. Compared to the control, vessel diameters highlighted using colored markers (diamond, hexagon and star) increased on the left side between EDD4 and EDD6 for all shapes, while the right side showed a decreasing trend **(Figure 4B)**. Additionally, the vessel diameters were largest in the triangle shape, intermediate in the circle, and smallest in the square. While the left-right symmetry of the body axis is well studied^[27]^, the vascular symmetry in developing chicks has not been extensively reported.

**Figure 4.**
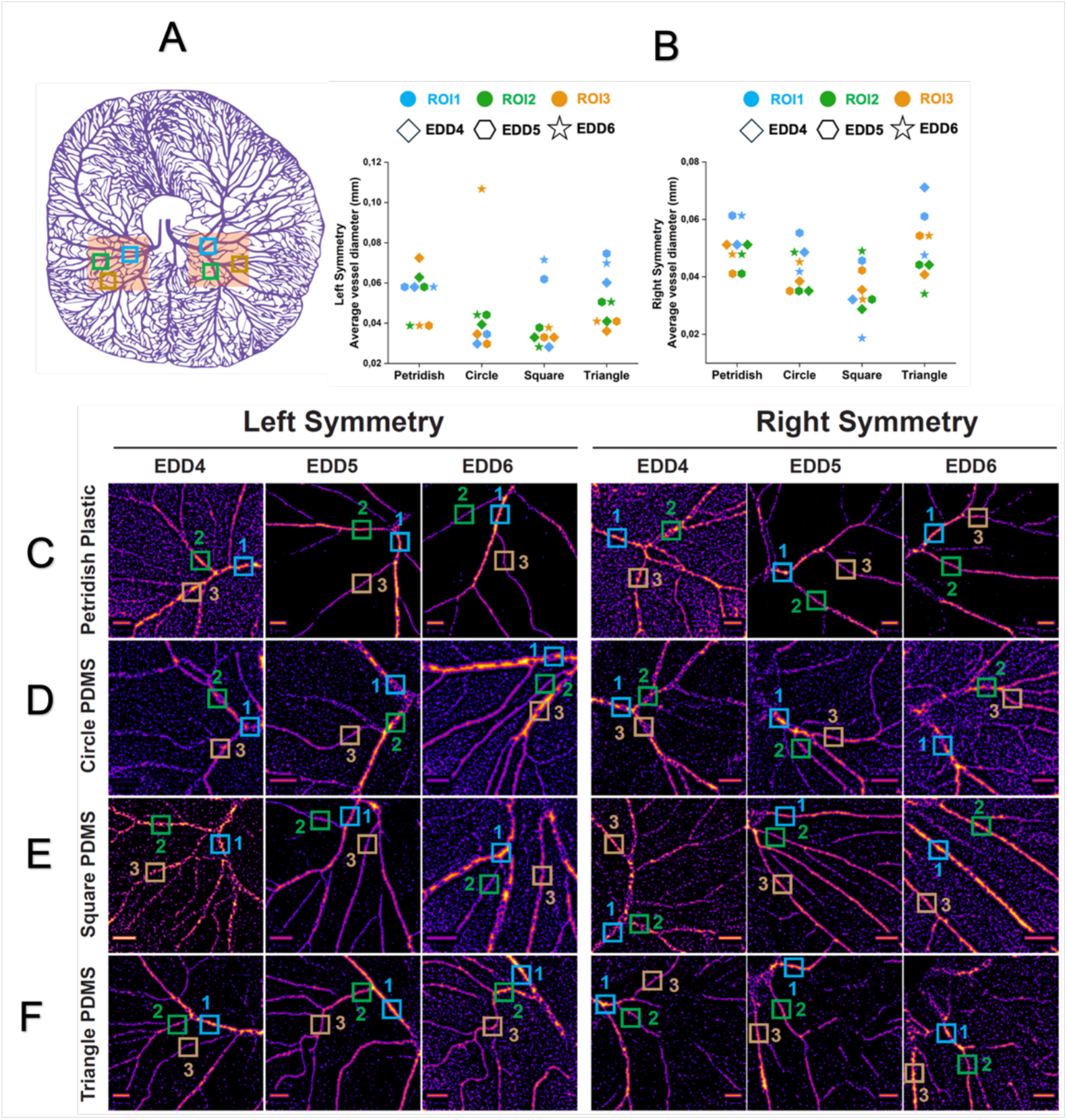
Effect of culture platform shape on left-right symmetry of developing vasculature. **(A)**represents the schematic mask of chick vascular network created using Adobe illustrator with the highlighted regions that were used for the quantitative analysis. **(B)** shows the graphical representation of quantitative data analysis results, compared with different culture platform and development days as indicated. Note the left and right symmetry regions highlighted in panel A have 3 selected data points (ROI) indicated via blue, green, brown color boxes, extracted from the multiple locations such as main branch, first and second hierarchical branches as indicated using with a pixel size of 25 × 25 respectively. Temporal information on the vessel diameters between EDD4 and EDD6 are represented using colored markers such as diamond, hexagon and star shapes respectively. **(C-F)** represents the spatial geometry to distance map highlighting the changes in vessel diameter. Measured values are calculated from 1 individual samples (n=1) and represented as mean ± SD along with individual data points. Statistical significance test was performed using OriginPro V.9.85 performing one way ANOVA and differences were considered statistically significant with P<0.05

To focus more on why the shape of the engineered eggshell platform affects the organization of the CAM vasculature, we compared changes in the shape of the yolk and the vascularized CAM in the different eggshell platforms over time. As the CAM can be seen as a thin membrane surrounding the yolk which has properties similar to a Newtonian liquid with a high viscosity^[28]^, shape changes in the yolk will result in distinct stress fields within the CAM. The circularity of the yolk and CAM is thus a critical indicator of the shape changes observed in this study, calculated using this formula C= 4π∗π*r*^2^/2πr. We compared changes in the shape of the yolk and CAM with the ideal shapes of a circle (C=1), square (C=0,785), and triangle (C=0,605) as earlier reported^[29]^. At EDD3, the transfer of egg contents to the eggshells resulted in an initial circularity value of the yolk and CAM of circular shape (C_yolk_=0,952 and C_CAM_=0,946), square shape (C_yolk_ = 0,955 and C_CAM_=0,894), and the triangular shape (C_yolk_ = 0,957 and C_CAM_=0,875) as shown in **Figure 5B**. Over development, at EDD6, significant changes in the circularity of both yolk and CAM were observed, resulting in circular shape (C_yolk_=0,891 and C_CAM_=0,912), square shape (C_yolk_ = 0,87 and C_CAM_=0,901) and triangular shape (C_yolk_ = 0,665 and C_CAM_=0,66) respectively. These shape changes in yolk and CAM greatly influenced the vessel orientation as demonstrated in **Figure 5C**. A more random organization of vessels was observed in Petridish and circle shapes, whereas a directional organization was seen in the square and triangle shape.

**Figure 5.**
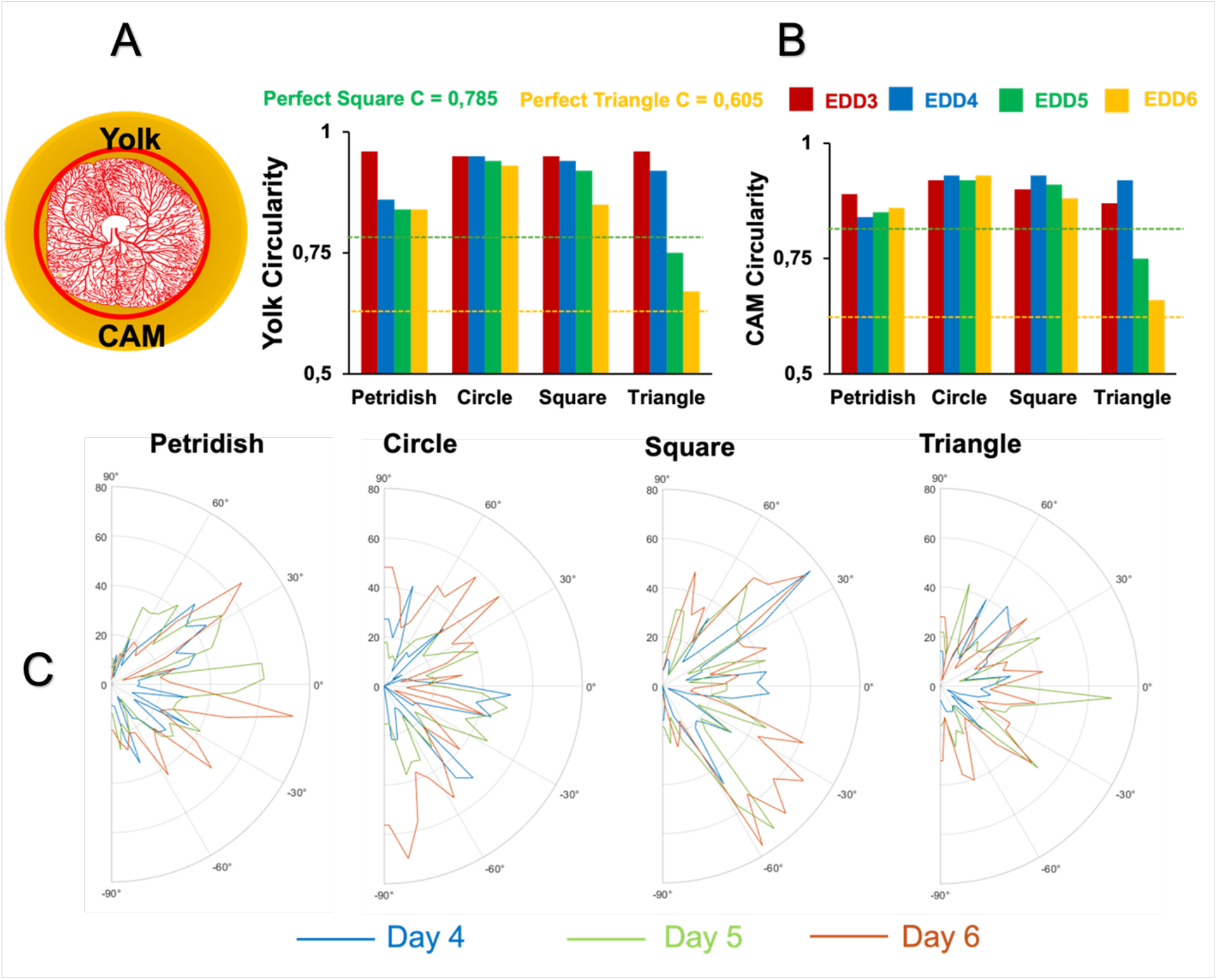
Effect of culture platform shape on yolk and CAM circularity and vasculature orientation. **(A)** represents the schematic mask of chick vascular network created using Adobe illustrator with highlighted regions of interest (ROI) represented for the quantitative analysis. Panel (B) show the mean circularity values of yolk and CAM respectively for the different culture platform. (C) represents the orientation maps of vessel structures for different development days for different culture platform. Statistical significance test was performed using OriginPro V.9.85 performing one way ANOVA and differences were considered statistically significant with P<0.05

### 3.4. Shape dynamics affects local and global vascular organization

Considering that every tissue, whether native or engineered, undergoes local deformation over time, we further investigated whether changes within the shape of the yolk influence vascular organization locally and globally. For this purpose, we fabricated a shape-shifting device as described in methods section 2.3. We selected the circle shape as the starting point as it closely resembles the shape of a natural egg. Figure 6B shows that before actuation of the device, the entire yolk and CAM vasculature from EDD4 adopt this circular shape. By applying compression actuation resulting in the change of the shape towards a square for one hour, we observed interesting changes in the vessel morphometrics at five different locations within the CAM vasculature (ROIs highlighted with colored circles).

**Figure 6.**
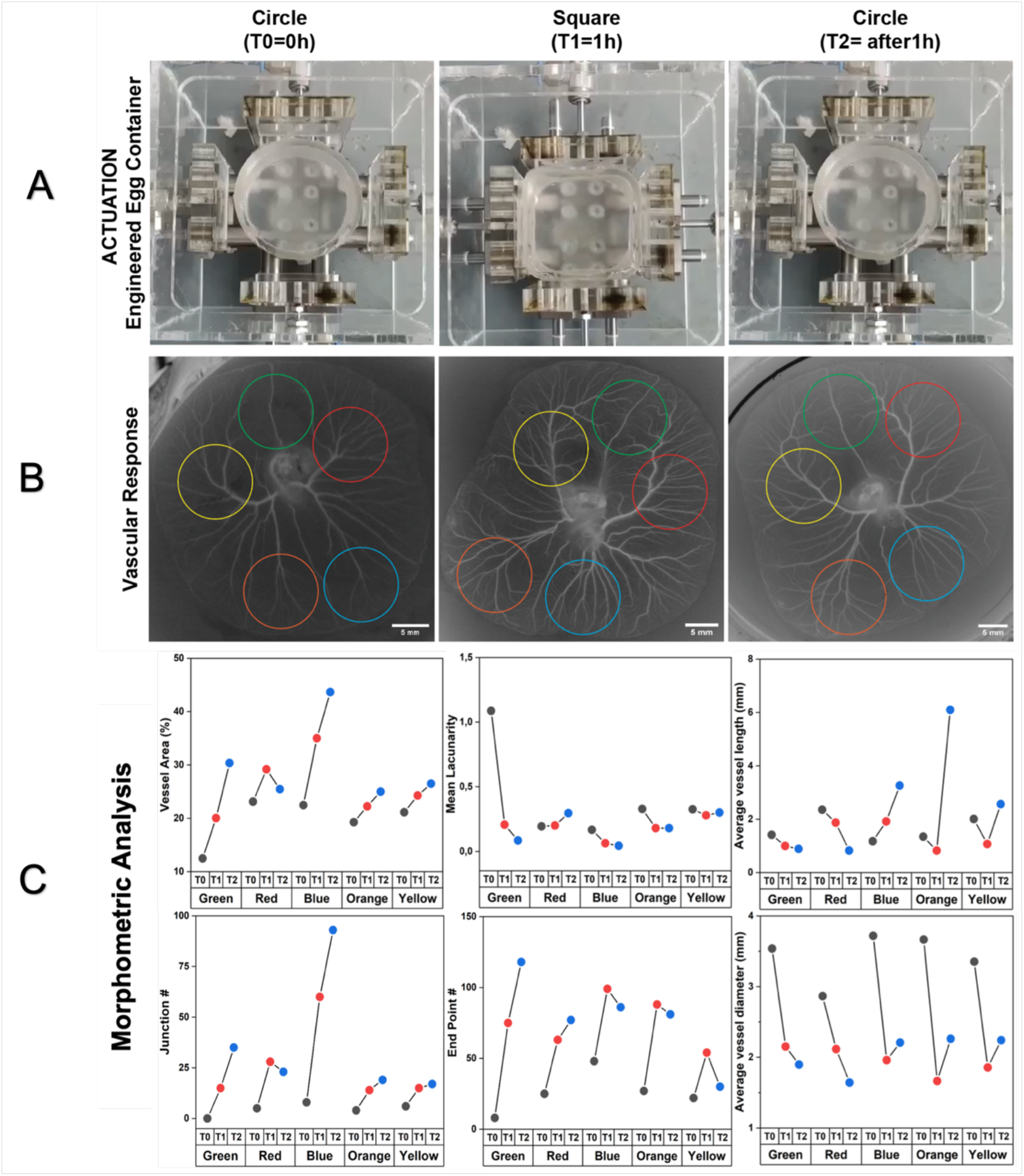
Shape-shifting culture platform influences vascular organization spatially. Panel (A) represents the actuation position of the shape-shifting culture platform from circle to square to circle. Panel (B) shows the snapshots of respective vascular response to the actuation of shape change. Note the highlighted color-coded ROIs were used for the morphometric evaluation. Panel (C) show the graphical representation of quantitative data analysis results, vessel morphometrics such as area, lacunarity, junction #, end point #, vessel length and diameter. Measured values are calculated from 1 individual sample (n=1) and represented as scatter plot with connected lines, with individual data points.

To ensure that this shape manipulation exclusively impacts remodelling rather than developmental growth, we examined vessel morphometrics at these locations across three time points; before the shape change is induced (T0), one hour after the shape has been changed towards a square (T1) and one hour after the shape reverted again to a circle (T2). We observed a consistent increase in vessel area over time. However, overall lacunarity showed a decreasing trend. These observations indicate that developmental growth is normal, as we previously reported^[25]^. Interestingly, at one specific location (red ROI), we noted a contrary pattern: a decrease in vessel area accompanied by an increase in lacunarity.

We subsequently analyzed the average vessel length and diameter across these locations. Interestingly, a consistent trend was observed across all regions, though significant differences were noted between the green and red regions compared to the blue, orange, and yellow regions. Previously, we observed that from EDD3 to EDD6, there was a similar increasing trend in diameter and length, indicating that vascular remodeling becomes more multi-scale over time^[25]^. In comparison to our previously reported values at EDD4, the current data demonstrate higher morphometric values, attributed to enhanced perfusion within the vessels induced by shape shifting, as illustrated in **Supplementary Video S5 and S6**.

Lastly, we compared the junction / branching number with the endpoint number across these locations. Here, we identified opposite trends between branching and endpoint numbers. There was an increasing trend in branching after actuation (except in the red ROI), while a decreasing trend was observed in endpoint numbers after actuation (except in the green and red ROI). A more random organization of vessels was observed before actuation, then it becomes directional organization after the actuation as shown in **Supplementary Figure S7**. As previous reported investigations have shown that in vivo mechanical stretching can alter tissue axis^[30]^, angiogenic sprouting^[31]^, vascular patterning^[8,9,12,13,32]^ and organoid geometric control ^[33]^, we anticipate that this study can contribute to the future investigations involving the development of shape shifting tissues.

In conclusion, our study demonstrates that the shape of eggshells housing chick vascularized CAM tissues influences their spatial and temporal organization during development. We attribute these alterations to the deformation of the yolk induced by the corners of the eggshell. To the best of our knowledge, this is the first investigation to examine how both static and dynamic shape changes influence the formation and organization of vasculature within a live organism. While previous studies have examined various shapes in vitro, the shapes we tested here do not replicate the complexity of native vascularized organs. However, these simple shapes are effective for observing significant vasculature morphological changes both at local and global spatial scales within the developing organism. This type of experiments has the potential to open new research avenues and inspire future investigations suggesting the incorporation of complex shapes into bioengineered tissues.

## Acknowledgements

The authors acknowledge the financial support from the ERC consolidator grant (724469) and NWO XS grant (OCENW.XS. 021) of Jeroen Rouwkema. Illustrations were made using Adobe Illustrator, and Biorender. The authors thank the Design Lab of the University of Twente for the fabrication of shape shifting prototypes.

## Author Contributions

PP and JR conceived the study. NSM and JR acquired the funding. PP fabricated culture platforms of different shapes, performed experiments, analyzed the data and wrote the original draft manuscript. PP and DW designed and fabricated the shape shifting prototype. OKK performed the analysis of orientation maps. All authors participated in the review and editing of the final version of the manuscript.

## Supplementary Information

### S1. Transfer of chick egg contents to PDMS based egg container

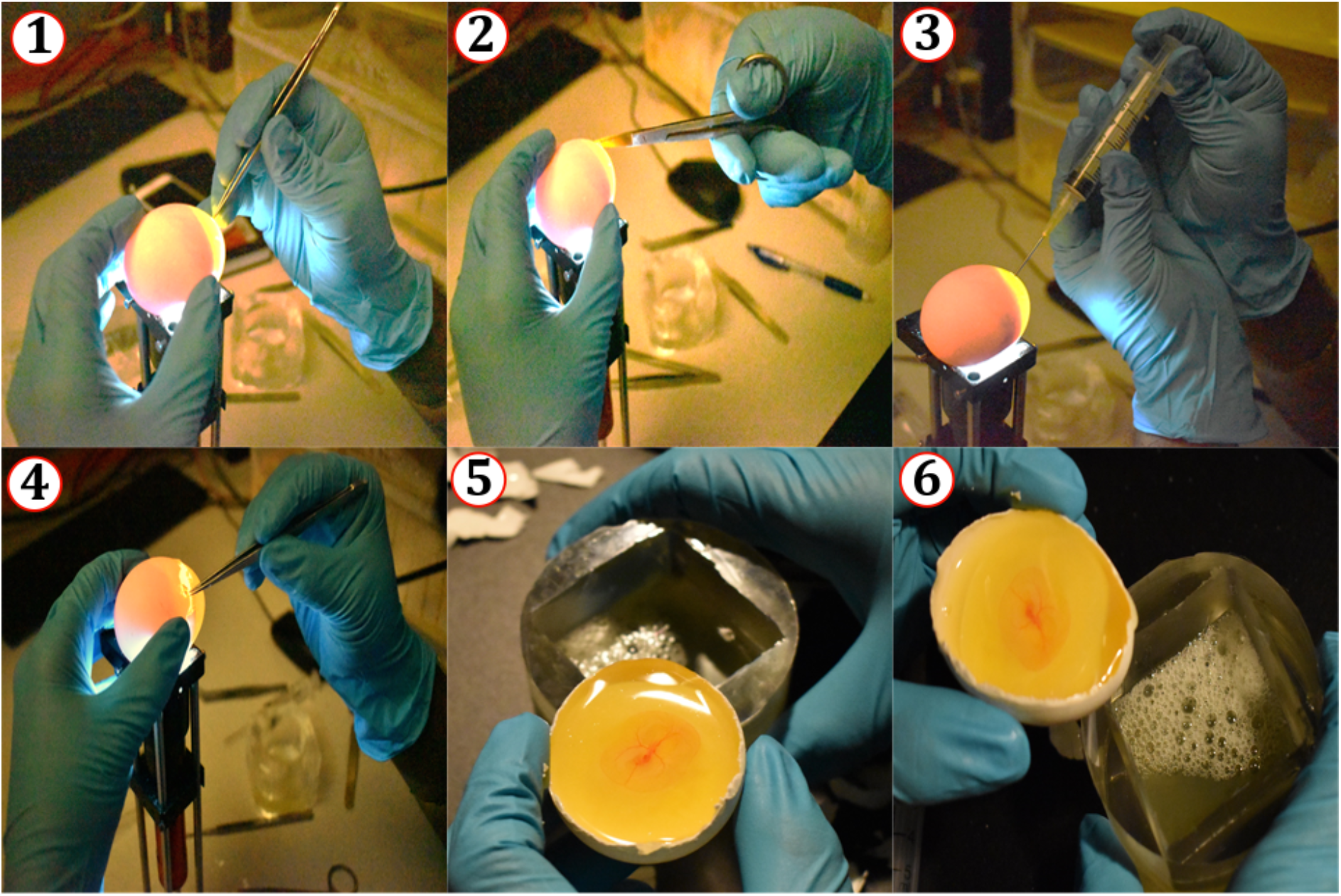
**(1)** Creation of 2mm hole **(2)** puncturing the outer eggshell layer **(3)** removal of albumen **(4)** cracking the eggshell **(5)** albumen transfer and positioning the embryo in the middle and **(6)** transfer to egg container

### S2. Fabrication of shaped engineered egg containers

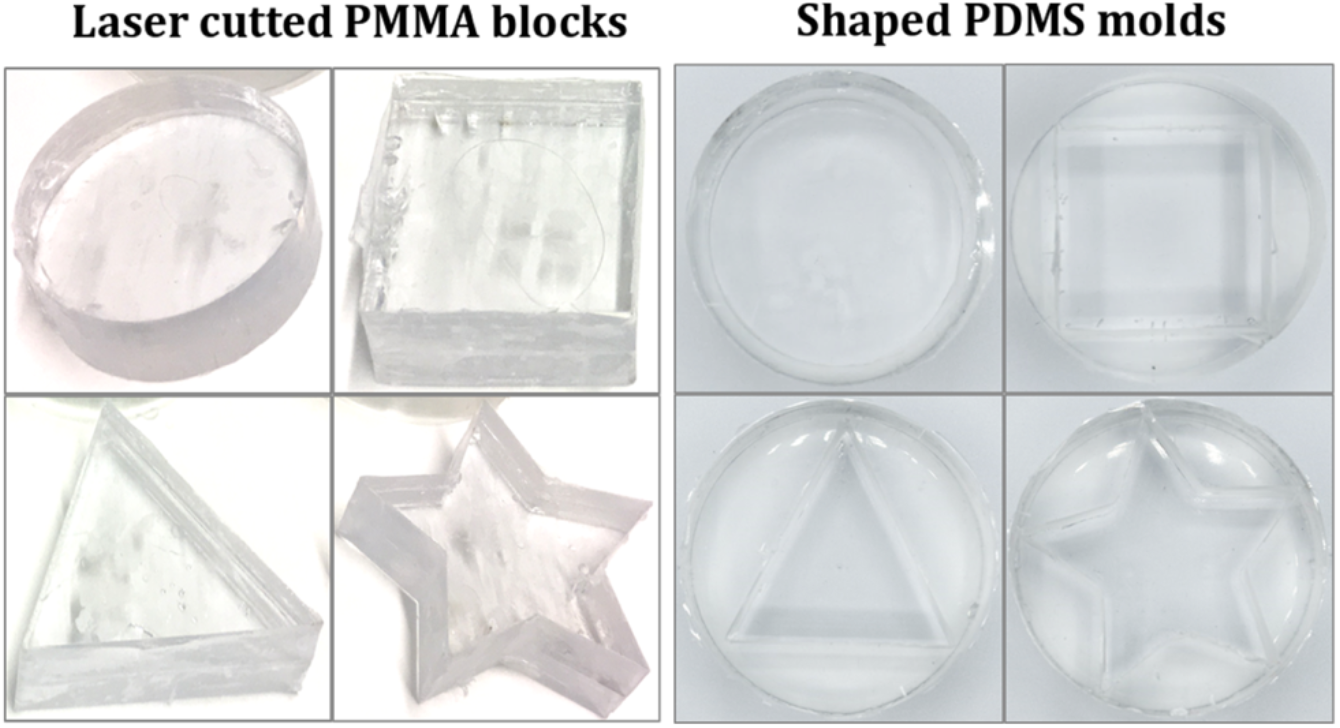

### S3. Chick CAM development within star-shape container

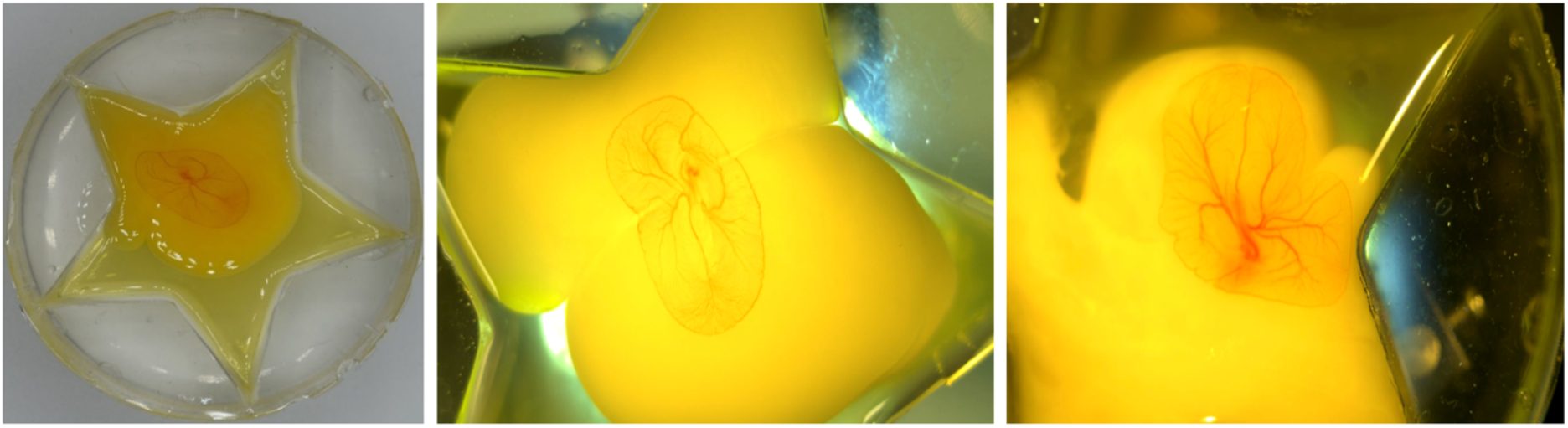

### S4. Vessel morphometrics analysis pipeline

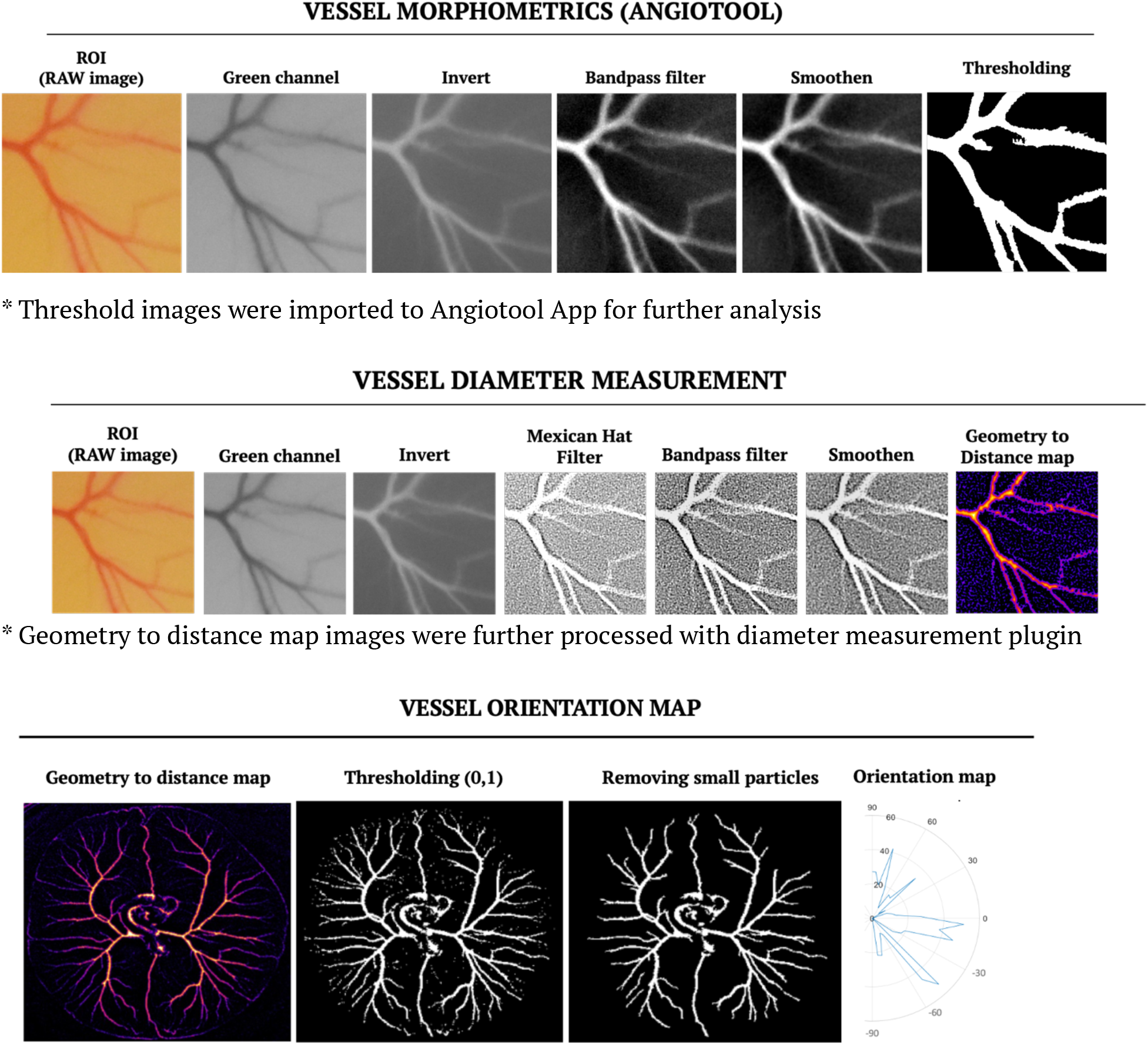

### S5. Shape influences vessel diameter

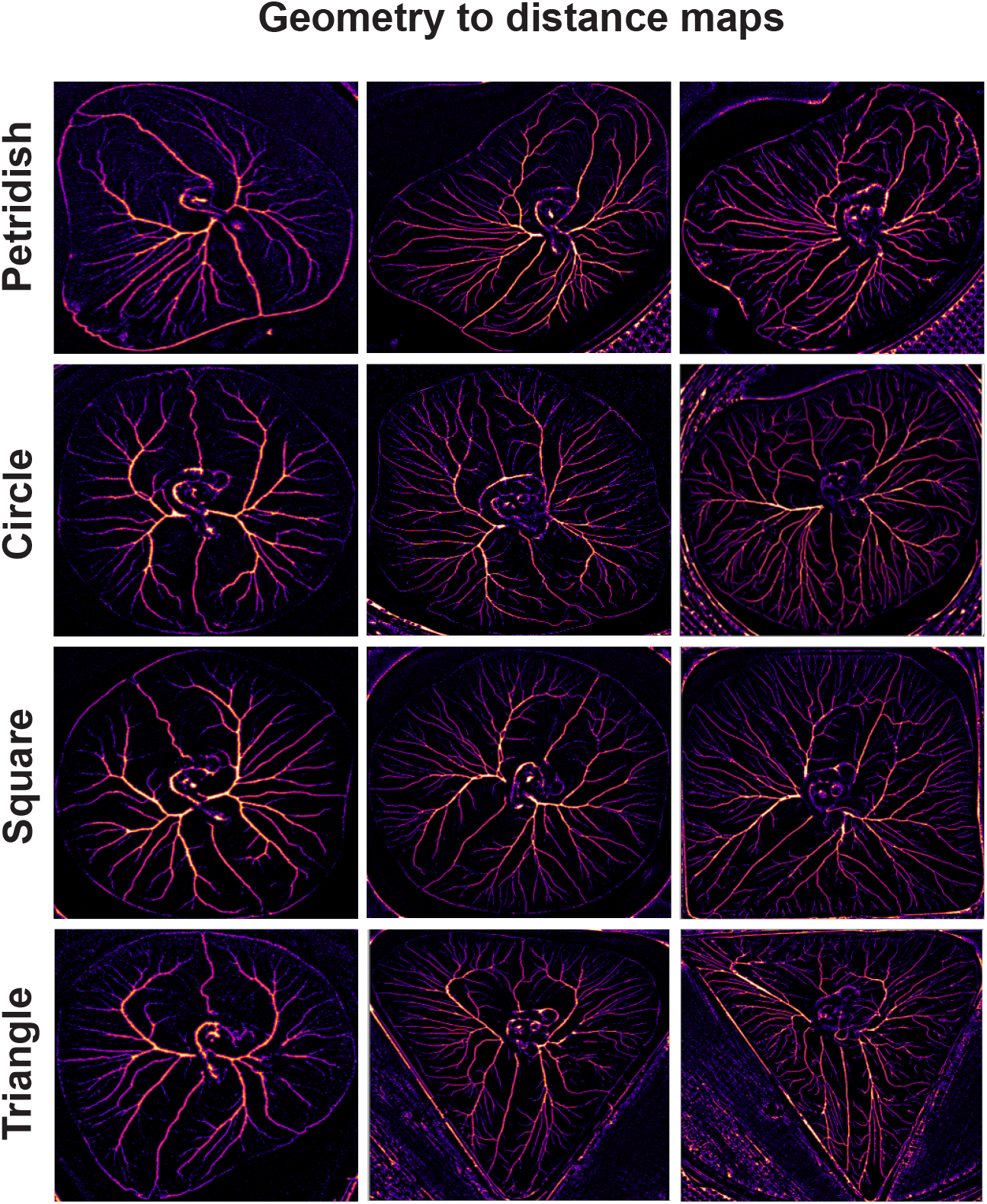

### S6. Vascular organization of chick CAM before and after shape change

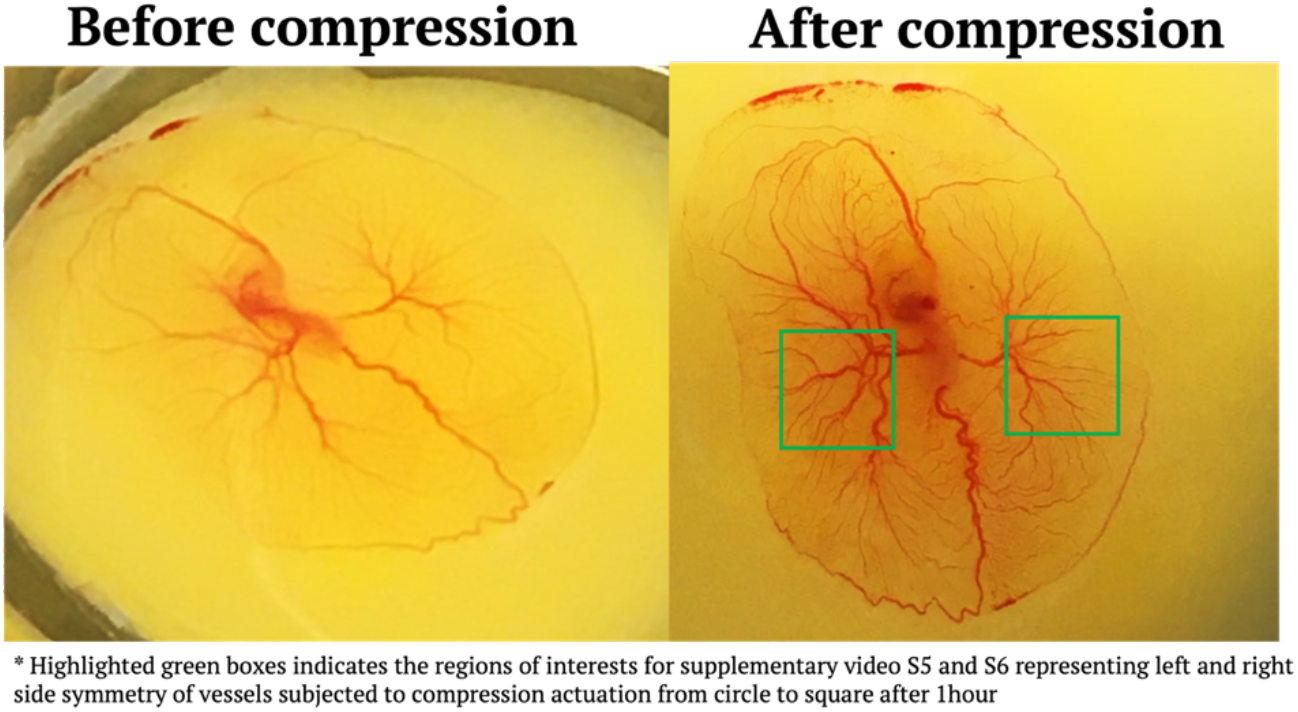

### S7. Shape-shifting influences vascular orientation

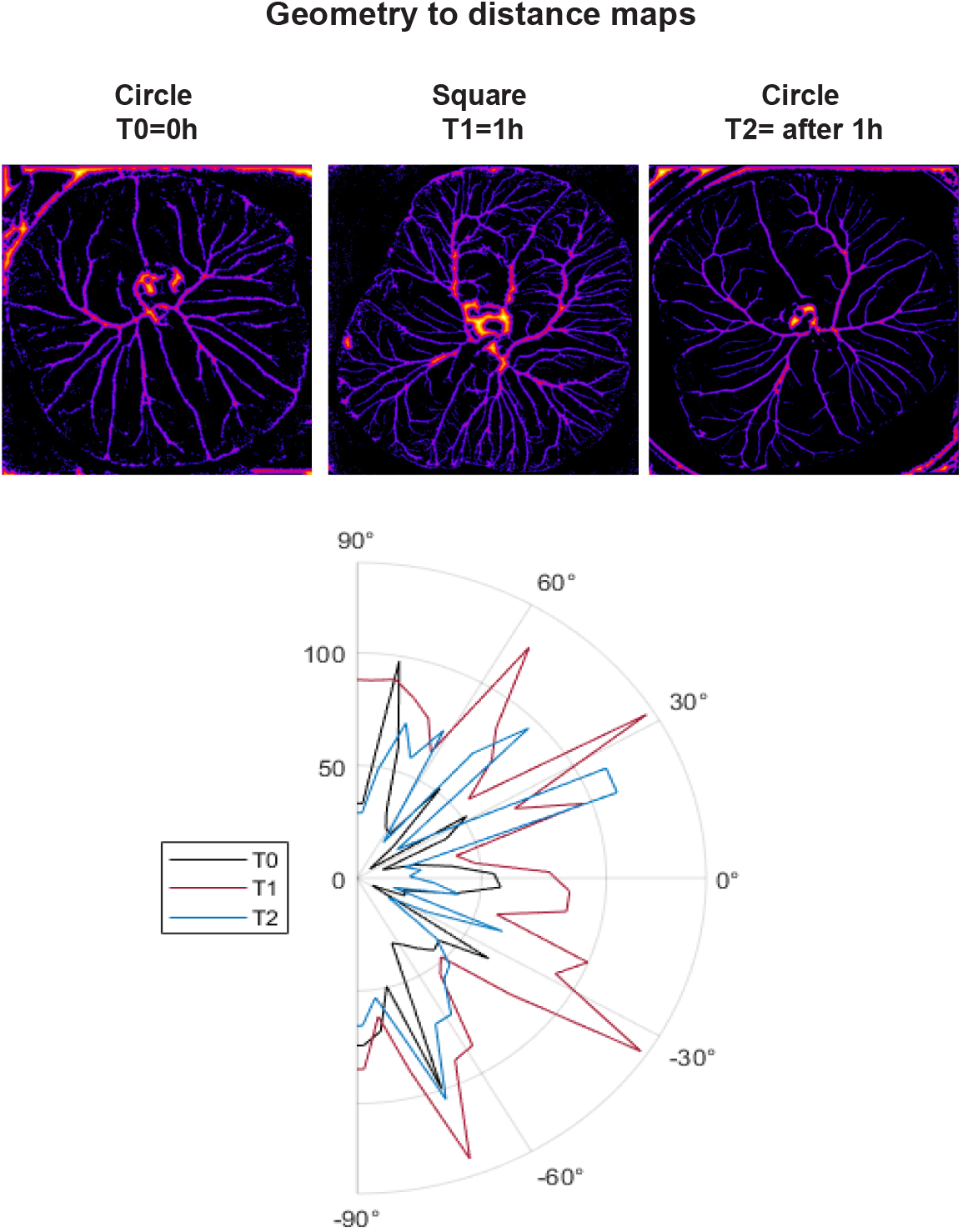

## Supplementary Video

Supplementary Video 1 Chick CAM in star shape container day 3 of development

Supplementary Video 2 Chick CAM in star shape container day 4 of development

Supplementary Video 3 Shape shifting prototype actuation of the PDMS egg container

Supplementary Video 4 Shape shifting prototype actuation with chick CAM

Supplementary Video 5 Vessel perfusion in left side symmetry CAM after actuation

Supplementary Video 6 Vessel perfusion in right side symmetry CAM after actuation

