## Supplementary figures and images for "Shape that matters: Yolk geometry spatially modulates developing vascular networks within chick chorioallantoic membrane"

### Supplementary Video S1

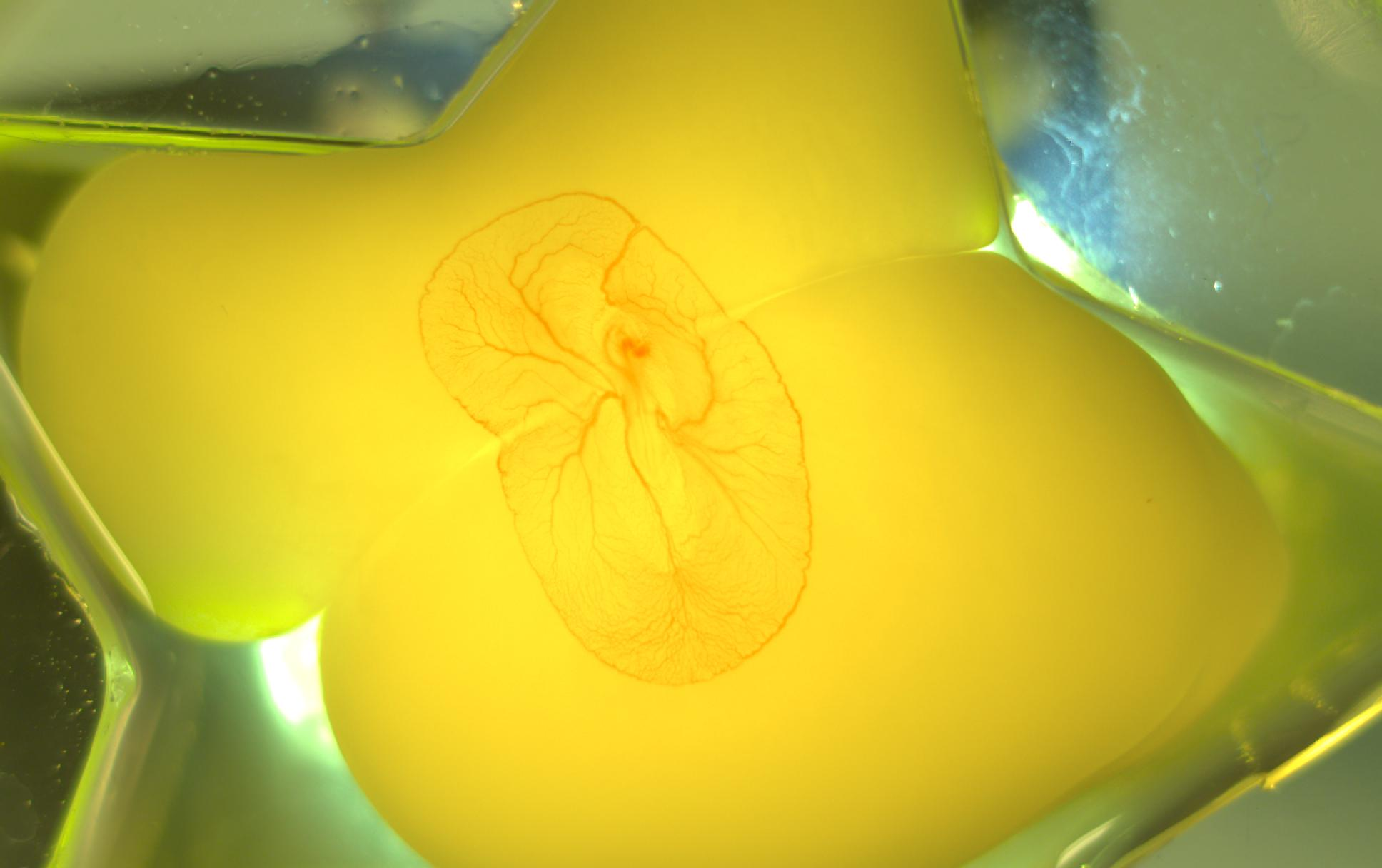

### Supplementary Video S2

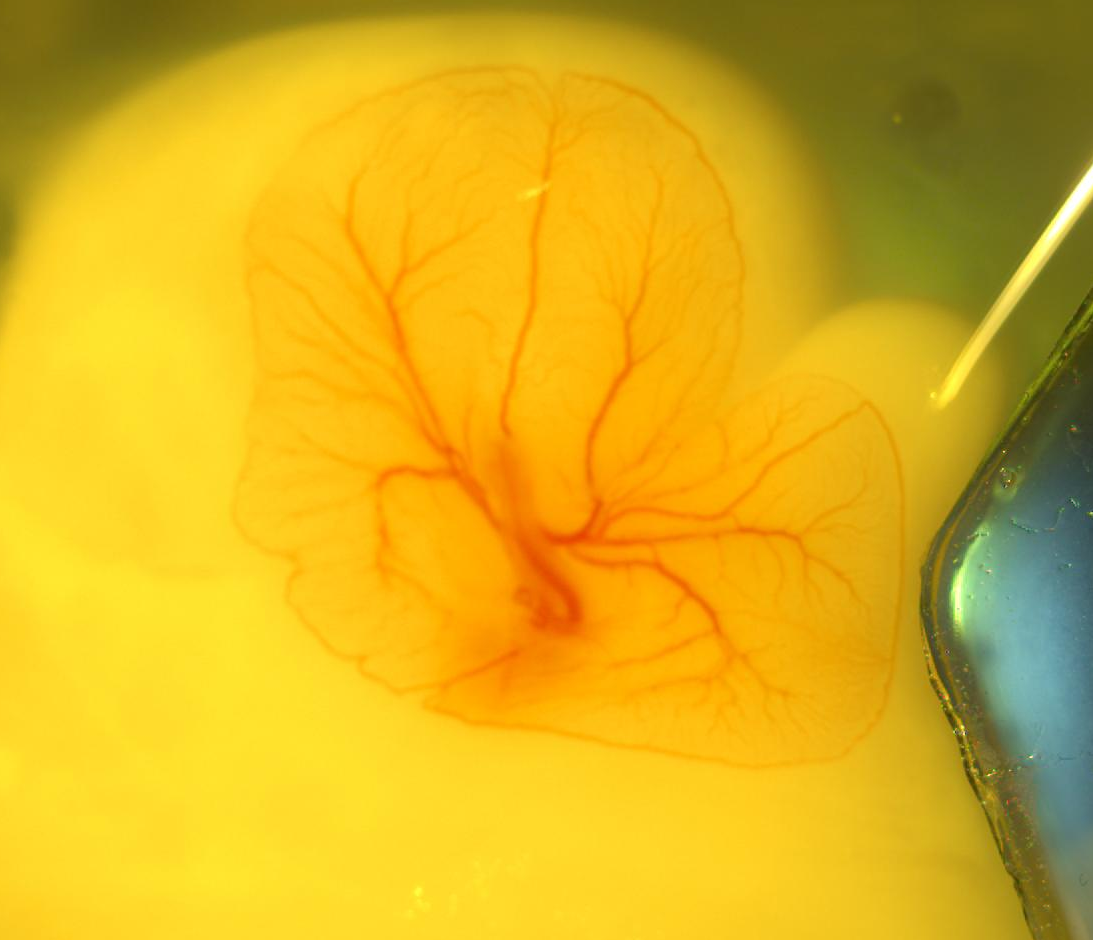
